# Disentangling neural correlates of tinnitus and hyperacusis following noise exposure in auditory cortex of rats

**DOI:** 10.1101/2024.05.01.592128

**Authors:** Naoki Wake, Tomoyo I. Shiramatsu, Hirokazu Takahashi

## Abstract

Both tinnitus and hyperacusis, likely triggered by hearing loss, can be attributed to maladaptive plasticity in auditory perception. However, owing to their co-occurrence, disentangling their neural mechanisms proves difficult. We hypothesized that the neural correlates of tinnitus are associated with neural activities triggered by low-intensity tones, while hyperacusis is linked to responses to moderate- and high-intensity tones. To test these hypotheses, we conducted behavioral and electrophysiological experiments in rats 2 to 8 days after traumatic tone exposure. In the behavioral experiments, prepulse and gap inhibition tended to exhibit different frequency characteristics (although not reaching sufficient statistical levels), suggesting that exposure to traumatic tones resulted in hyperacusis and tinnitus symptoms at different frequency ranges. When examining the auditory cortex at the thalamocortical recipient layer, we observed that tinnitus symptoms correlated with a disorganized tonotopic map, typically characterized by responses to low-intensity tones. Neural correlates of hyperacusis were found in the cortical recruitment function at the multi-unit activity (MUA) level, but not at the local field potential (LFP) level, in response to moderate- and high-intensity tones. This shift from LFP to MUA was associated with a loss of monotonicity, suggesting a crucial role for inhibitory synapses. Thus, in acute symptoms of traumatic tone exposure, our experiments successfully disentangled the neural correlates of tinnitus and hyperacusis at the thalamocortical recipient layer of the auditory cortex. They also suggested that tinnitus is linked to central noise, whereas hyperacusis is associated with aberrant gain control. Further interactions between animal experiments and clinical studies will offer insights into neural mechanisms, diagnosis and treatments of tinnitus and hyperacusis, specifically in terms of long-term plasticity of chronic symptoms.

## 1 Introduction

Hearing loss is often associated with a high comorbidity of tinnitus and hyperacusis. For instance, more than 60% of patients with tinnitus also experience hyperacusis, and conversely, over 80% of patients with hyperacusis suffer from chronic tinnitus (Anari et al., 1999; Dauman and Bouscau-Faure, 2005; Schecklmann et al., 2014b; Sztuka et al., 2010). Both tinnitus and hyperacusis likely result from maladaptive changes in gain control within the auditory system. This maladaptive gain control is characterized by an increase in the activity of the central auditory pathway, which is triggered by a decrease in peripheral input (Auerbach and Gritton, 2022; Auerbach et al., 2019; Auerbach et al., 2014; Eggermont and Roberts, 2004; Herrmann and Butler, 2021; Roberts et al., 2010; Robinson and McAlpine, 2009; Schaette and Kempter, 2006; Shore et al., 2016; Zeng, 2013). The maladaptive gain control may be attributed to homeostatic plasticity, a mechanism that maintains baseline activity levels following perturbations (Turrigiano, 2012; Turrigiano and Nelson, 2004). In the context of hearing loss, this homeostatic plasticity can distort neural representations and lead to significant auditory perceptual challenges (Eggermont, J. J., 2017; Herrmann and Butler; Nahmani and Turrigiano, 2014; Norena, 2011; Schaette and Kempter, 2006). This distortion can involve synaptic sensitization through receptor upregulation (Balaram et al., 2019; Sturm et al., 2017), synaptic disinhibition (Balaram et al., 2019; Sanes and Kotak, 2011; Sarro et al., 2008; Sturm et al., 2017), and increased intrinsic excitability (burstiness) (Li et al., 2013; Li et al., 2015; Pilati et al., 2012; Shore et al., 2016; Wu et al., 2016; Yang et al., 2012). Additionally, non-homeostatic regulation of sound intensity may also play a role, as gain control following acoustic trauma is heterogeneous among pyramidal neurons in the auditory cortex (McGill et al., 2022).

Acoustic trauma reduces spontaneous and sound-evoked activities at the auditory nerve (Heinz et al., 2005; Heinz and Young, 2004; Hickox and Liberman, 2014; Kiang et al., 1970; Wake et al., 1993; Wang et al., 1997). However, in response to this trauma, it induces hyperactivity and synchrony in spontaneous and sound-evoked activities at the cochlear nucleus (Cai et al., 2009; Kaltenbach and Afman, 2000; Kaltenbach and McCASLIN, 1996; Kaltenbach et al., 2000; Wu et al., 2016), the inferior colliculus (Bauer et al., 2008; Hesse et al., 2016; Hickox and Liberman, 2014; Mulders and Robertson, 2009; Salvi et al., 1990; Sun et al., 2011; Xiong et al., 2017), and the auditory cortex (Asokan et al., 2018; Basura et al., 2015; Chambers et al., 2016; Komiya and Eggermont, 2000; Kotak et al., 2005; Norena and Eggermont, 2003; Norena et al., 2003; Parameshwarappa et al., 2022; Popelar et al., 1987; Qiu et al., 2000; Resnik and Polley, 2017; Resnik and Polley, 2021; Seki and Eggermont, 2003; Sun et al., 2012; Syka et al., 1994; Wong et al., 2020; Yang et al., 2011). Furthermore, resting-state fMRI in rats with drug-induced depression of the cochlea has revealed hyperactivity in various brain regions, including the cerebellum, reticular formation, amygdala, hippocampus, and the higher-order auditory pathway, which encompasses the inferior colliculus, medial geniculate body, and auditory cortex (Chen et al., 2015).

The aberrant neural activities are believed to underlie hyperacusis, defined as “a reduced tolerance to sounds that are perceived as normal by the majority of the population” (Adams et al., 2021). In hyperacusis, moderate-intensity sounds are perceived as intolerably loud, aversive, or even painful (Anari et al., 1999; Auerbach et al., 2019; Auerbach et al., 2014; Baguley, 2003; Eggermont, J. J., 2017; Pienkowski, 2017, 2019). Human imaging studies further support the presence of sound-evoked hyperactivity across multiple auditory nuclei in patients with hyperacusis (Bigras et al.; Gu et al., 2010; Hwang et al., 2009; Koops and van Dijk, 2021; Lanting et al., 2008; Lockwood et al., 1998; Melcher et al., 2009). This hyperactivity is also associated with a steep growth function of sound-evoked activities concerning test intensity (Auerbach et al., 2019; Zeng, 2013, 2020).

While hyperacusis arises from abnormal gain of evoked responses, one of possible mechanisms of tinnitus is increased central noise, independent of gain (Gu et al., 2010; Knipper et al., 2013; Zeng, 2013) (but some studies note that changes in firing rates are not necessarily tinnitus specific (Coomber et al., 2014; Longenecker and Galazyuk, 2016; Ropp et al., 2014)). This model predicts gain reduction (Hofmeier et al., 2018) and a dissociation between cortical and subcortical activities (Boyen et al., 2014) in tinnitus frequency. Conversely, it predicts gain increase (Diehl and Schaette, 2015) and hyperactivity in the frequency related to hyperacusis, affecting both subcortical and cortical activities (Chen et al., 2015; Gu et al., 2010; Knipper et al., 2013; Ruttiger et al., 2013). However, it can be challenging to disentangle these effects, primarily owing to the co-occurrence of tinnitus and hyperacusis (Cederroth et al., 2020; Lanting et al., 2008; Melcher et al., 2009; Schecklmann et al., 2014a). Consequently, hyperactivity within a specific region on the tonotopic map in the auditory cortex is considered a neural signature of either tinnitus or hyperacusis (Auerbach et al., 2019; Herrmann and Butler, 2021; McGill et al., 2022; Norena, 2011), which has not been reliably distinguished based on their associated symptoms.

In this study, we aimed to disentangle the neural correlates of tinnitus and hyperacusis within the auditory cortex of noise-exposed rats. Several behavioral tests have been employed to estimate tinnitus and hyperacusis in animal models (Hayes et al., 2014). Specifically, prepulse and gap inhibitions (PPI and GPI) of acoustic startles have been established as behavioral indicators of hyperacusis and tinnitus symptoms in animals, respectively, in a similar fashion of experiments (Turner and Larsen, 2016). Furthermore, despite being a reflexive measure, we recently demonstrated that PPI can predict subjective pure-tone audiometry based on operant conditioning in rats (Wake et al., 2021).

Our initial focus was to confirm the distinct characteristics of PPI and GPI in rats with noise-induced hearing loss, which suggest that acoustic trauma made hyperacusis and tinnitus in different frequencies. Subsequently, we delved into the neural correlates of PPI (i.e., hyperacusis symptoms) and GPI (tinnitus) within the auditory cortex using high-density microelectrode array mapping. Given that tinnitus is associated with increased central noise, we expected to find the neural correlates of tinnitus in response to faint tones with a low signal-to-noise (S/N) ratio. Accordingly, we hypothesized that GPI is correlated with the extent of spatial disorganization of the characteristic frequency (CF) in the auditory cortex, which is typically characterized with low-intensity tones. Furthermore, we postulated that, unlike the putative neural correlates of tinnitus, hyperacusis-like symptoms are associated with neural gain from synaptic inputs to spike outputs in response to moderate- and high-intensity tones.

To investigate this, we measured neural activities at both the local field potential (LFP) and multi-unit activity (MUA) levels, and attempted to characterize the neural gain from LFP to MUA. We here assumed that the first negative deflection of auditory-evoked LFPs in layer 4 reflected synaptic inputs of thalamo-cortical projection rather than suprathreshold discharges, based on previous physiological and simulation studies (Einevoll et al., 2013; Haider et al., 2016; Mazzoni et al., 2015). Assuming that LFP and MUA represent cortical inputs and cortical responses, respectively, our hypothesis predicts that the neural gain from LFP to MUA is correlated with the enhancement of PPI induced by trauma. To estimate the neural gain, we quantified the cortical recruitment, or the activation level of the entire auditory cortex, from either tone-evoked LFP or MUA. We attempted to compare the cortical recruitment between LFP and MUA to investigate how efficiently the thalamo-cortical synaptic inputs were converted into cortical discharges.

## 2 Materials and Methods

This study was conducted in accordance with the National Institutes of Health guide for the care and use of Laboratory animals (NIH Publications No. 8023, revised 1978) and following the recommendations of the ARRIVE guidelines (https://arriveguidelines.org/). All the procedures that involved the care and use of animals were approved by the Committee on the Ethics of Animal Experiments at the Research Center for Advanced Science and Technology, The University of Tokyo (RAC170005). Surgery, traumatic noise exposure, and neural recording were performed under isoflurane anesthesia (3% for induction and 1–2% for maintenance), and every effort was made to minimize the suffering of the animals. All experiments were carried out in a sound-attenuating chamber (AMC-4015; O’Hara & Co. Ltd., Tokyo, Japan), where the background noise level was 32.1 dB (A-weighted equivalent continuous sound level; MT-325, Mothertool Co., Ltd, Nagano, Japan).

### 2.1 Animals

A total of 16 male Wistar rats (7–10 weeks old; body weight, 250-460g) were used in this study. In eight of the 16 rats, acoustic trauma was induced in the left ear by exposing them to a 10-kHz tone with an intensity of 125 dB SPL (Sound Pressure Level in dB with respect to 20 µPa) for one hour using a loudspeaker (Selenium ST 400, Los Angeles, CA) (Wake et al., 2019). To safeguard the hearing in the right ear during exposure, a silicone impression material (Dent Silicone-V, Shohu, Kyoto, Japan) was injected into the right ear canal. PPI and GPI in these exposed animals were recorded before and 2–8 days after the traumatic noise exposure to measure changes in their hearing sensitivity to tones. Immediately following the recording of both PPI and GPI for post-exposure measurements, LFP and MUA were recorded in the fourth layer of the right auditory cortex. The remaining eight animals were designated as a control group, and PPI, GPI, LFP, and MUA were recorded without exposing them to noise.

### 2.2 Behavioral experiments

The procedures for measuring PPI were described in detail in our previous work (Wake et al., 2021). Briefly, an acoustic startle stimulus presented through a speaker (DDL-RT16C, Alpine, Tokyo, Japan) was used as white noise (95-dB SPL, 10 ms). The startle reflex was measured using a force sensor attached to the floor where the animals were placed (PW6C 5KG; Unipulse Corp., Tokyo, Japan). The responses of the startle reflex were recorded using a DA converter (USB-6461, National Instruments, Austin, TX) and quantified by measuring the peak-to-peak magnitude of the force sensor output from 100 ms before to 200 ms after the startle stimulus emission. For each tone frequency (f [kHz]), which included both with and without prepulse conditions, PPI (and GPI) were defined as follows in Equation (1):

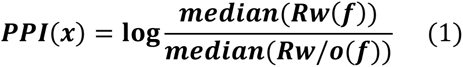

where Rw and Rw/o denote startle reflexes with and without the prepulse (and gap), respectively. Since the spontaneous movements of animals could sometimes interfere with the accurate detection of startle reflexes during certain trials (Tziridis et al., 2012), trials where the startle reflexes exceeded 2σ of the mean were excluded. The same formula and criteria were also applied to GPI(f), where startle reflexes were measured both with and without gaps when background tones with different frequencies were presented. Both PPI(f) and GPI(f) yield positive values when a prepulse or gap inhibits the startle reflexes.

GPI was first measured with five distinct background sounds; four continuous pure tones (each at 60 dB SPL) with frequencies of 4, 8, 16, and 32 kHz, along with broadband noise (BBN; also at 60 dB SPL) spanning from 0.1 to 64 kHz. A 50-ms silent gap was introduced after 20±2 s of exposure to a background tone or noise. The time interval between the initiation of the startle stimulus and the gap period was set at 100 ms. At the beginning of GPI recording, rats were given a 120-s period to acclimate to the experiment environment. Following this, the startle stimuli were presented twice for habituation (Ison et al., 1973). The GPI recording encompassed 10 trials, each involving background sounds presented in a random order, both with and without gap conditions.

Subsequently, PPI was recorded after GPI. Five types of prepulses were used, including 50-ms tone bursts of 4, 8, 16, and 32 kHz (each at 60 dB SPL), along with a 50-ms BBN (60 dB SPL). The time interval between the startle stimulus and a prepulse was 100 ms. The startle stimulus was presented every 20±2 s. Similar to the GPI measurement, the PPI measurement started with a 120-s acclimatization period and two habituation stimuli, and consisted of 10 trials. Each trial tested the five prepulses in a random order, both with and without prepulse conditions. For PPI, seven of the exposed animals were analyzed, owing to missing PPI data for one subject.

### 2.3 Electrophysiology

The procedures for recording neural activity in the auditory cortex were described in detail in our previous work (Wake et al., 2019). Briefly, rats were anesthetized with isoflurane. Then, cisternal cerebrospinal fluid drainage was performed to avoid cerebral edema. After that, the auditory cortex was surgically exposed, and the dura mater over the auditory cortex was removed. Atropine sulfate (Abbott Japan Co. Ltd., Tokyo, Japan; 0.1 mg/kg) was administered at the beginning of the surgery to reduce the bronchial secretion viscosity, whereas xylocaine was subcutaneously administered for local anesthesia when necessary. After locating the auditory cortex through surface recording, a microelectrode array (ICS-96; Blackrock Microsystems, Salt Lake City, UT, USA) was inserted at the depth of 600 μm, specifically within the fourth layer, to record the auditory-evoked MUA and LFP in the auditory cortex. The array was equipped with 10×10 recording electrodes, featuring 400-μm inter-electrode spacing and covering an area of 4×4 mm2 that spanned the entire auditory cortex. The Cerebus Data Acquisition System (Cyberkinetics, Inc., Salt Lake City, UT, USA) amplified neural signals 1,000 times and recorded LFP and MUA. The filter passband and the sampling rates were set at 0.3–500 Hz and 1 kHz for LFP and 250–7500 Hz and 30 kHz for MUA, respectively. In online processing, multi-unit spikes were detected as threshold-crossing events with a threshold set at −5.65 times the root mean square from the average. No spike sorting was applied offline, assuming that MUA can be considered as spatial pooling of single unit activities as far as the number of units in MUA did not vary across recording sites. This assumption is reasonably valid in our low-impedance microelectrode recording (Noda and Takahashi, 2015).

The test stimuli consisted of tone bursts (5-ms rise/fall, 20-ms plateau) with varied frequencies and SPLs: 18 frequencies between 1.6 and 64 kHz with a 1/3-octave interval and seven SPLs from 20- to 80-dB SPL with a 10-dB interval. These test stimuli were calibrated with a 1/4-inch microphone at the pinna (4939, Brüel & Kjær, Nærum, Denmark) and were presented bilaterally through a speaker (DDLiner; Alpine Electronics of Australia Pty. Ltd., Hallam, Melbourne, VIC, Australia). For each combination of frequency and SPL, the test stimuli were presented 20 times in a pseudo-random fashion. The frequency response area (FRA) at each recording site was then determined by assessing the magnitude of neural responses as a function of tone frequency and SPL. The MUA magnitude was quantified as the number of tone-evoked spikes, defined as the difference between the total spike counts within 100 ms from the tone onset and the spontaneous spike counts within 4 ms from the tone onset (Guo et al., 2012; Noda and Takahashi, 2015). Additionally, LFP was characterized as the grand average across 20 trials, then the maximum amplitude (the first negative deflection) within the 10–60 ms time window was taken as the LFP magnitude (Takahashi et al., 2004, 2005a).

Based on the FRA, the CF at each recording site was determined as the test frequency where evoked MUAs were observed at the lowest SPL or where the largest evoked MUA was recorded at 20 dB SPL. The tonotopic maps within the auditory cortex were subsequently generated by spatially mapping these CF. Given that CF was characterized at the lowest possible SPL, these tonotopic maps were primarily characterized by low-intensity tones.

Additionally, cortical recruitment functions (CRFs) of tone-evoked MUA and LFP were also determined as a measure of the activation level of the entire auditory cortex for every condition of frequency and SPL (Kilgard and Merzenich, 1998; Takahashi et al., 2011). The CRFs were characterized across the entire range from the lowest SPL to the highest SPL tones. To construct the CRFs, the FRA was binarized based on a specified threshold, classifying each recording site as either active (1) or inactive (0) in response to each stimulus. The threshold to binarize the FRA was defined as previously described: For MUA, it was set as the inflection point of a smoothed z-score histogram of MUA (Guo et al., 2012; Noda and Takahashi, 2015); for LFP, it was determined as 1.2 times the baseline fluctuations within the 400–500 ms window after the stimulus onset (Liu et al., 2015). Using this binarized FRA (bFRA), CRF_MUA_ and CRF_LFP_ were defined as the proportion of active sites for a given stimulus, at the levels of MUA and LFP, respectively.

## 3 Results

### 3.1 Behavioral signature of hearing loss, hyperacusis, and tinnitus

In our behavioral experiments, based on PPI and GPI measurements taken before and after the traumatic noise exposure were used to interpret changes in hearing loss, tinnitus, and hyperacusis. A decrease in PPI, a decrease in GPI and an increase in PPI after noise exposure were considered indicators of hearing loss, tinnitus, and hyperacusis, respectively. After 1-h exposure to a 125-dB SPL, 10-kHz tone, a significant decrease in PPI to broadband noise was observed (pre- vs. post-exposure, p=0.0379, two-tailed Mann–Whitney U test), indicating the presence of noise-induced hearing loss (Fig. 1a). Although no significant changes were observed in PPI responses to individual pure tones, the difference in tone PPI (ΔPPI) between the pre- and post-noise exposure time points displayed a U-shaped pattern, with PPI decreases particularly evident in the 8–16 kHz range, suggesting a distinct hearing in this frequency range (Fig. 1b). Conversely, PPI values at 4 kHz and 32 kHz occasionally increased after noise exposure, implying that the noise-exposed animals became more sensitive to these specific tones than their unexposed counterparts. However, these signs of hyperacusis varied widely across subjects and did not reach statistical confirmation. The difference in GPI (ΔGPI) between pre- and post-exposure tended to decrease with the test frequency, and post-exposure GPI was marginally significantly smaller than pre-exposure GPI at 32 kHz (two-tailed Mann–Whitney U test, p = 0.0499; Fig. 1c, d). This trend suggested that the acoustic trauma induced tinnitus, particularly in the high-frequency range. No significant correlation was observed between ΔPPI and ΔGPI (Supplementary Fig. S1; R=0.231; t-test, p=0.238). Notably, the variance of ΔPPI tended to be larger than that of ΔGPI (F-test, p = 0.0458), indicating that hyperacusis symptoms exhibited more substantial variations across subjects compared to tinnitus symptoms.

**Figure 1.**
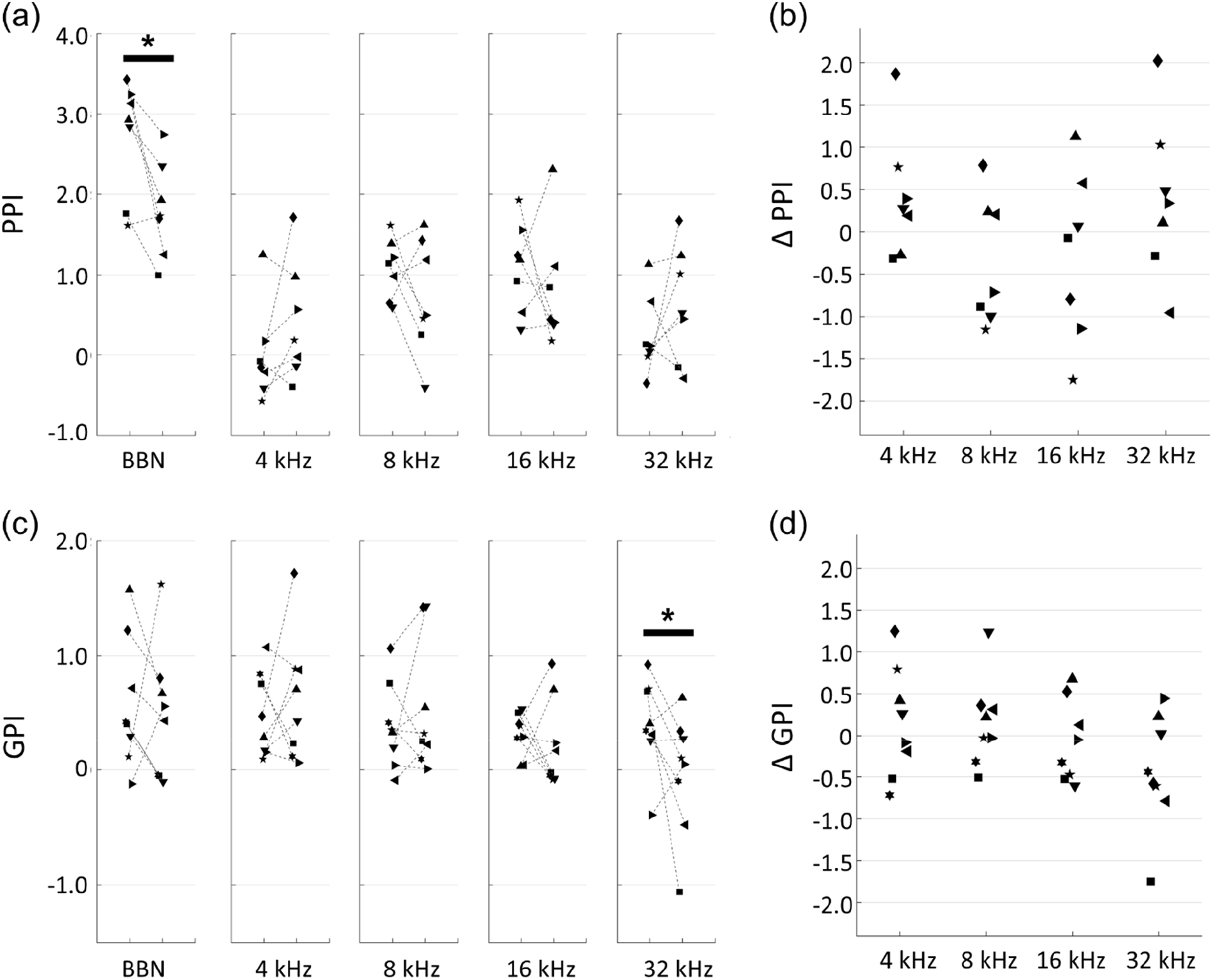
Behavioral experiments. (a) Prepulse inhibition (PPI) of pre- and post-exposure conditions. Broadband noise (BBN) and tones with indicated frequencies were used for prepulse. Each symbol indicates a different animal. Asterisks represent statistical differences between groups (p<0.05, Mann–Whitney U test). (b) PPI differences (ΔPPI) between pre- and post-treatment of acoustic trauma. (c) Gap-inhibition (GPI) of pre- and post-exposure conditions. (d) GPI differences (ΔGPI) between pre- and post-exposure.

### 3.2 Map disorganization

In the neural activity characterization, recording sites with CFs were considered to be within the auditory cortex and were used in subsequent analyses. Figure 2a displays representative CF maps in both the control and noise-exposed groups (see also Fig. 3a for the post-stimulus histogram of MUA for each test stimulus, which was used to determine CF). Consistent with previous studies (Funamizu et al., 2013; Shiramatsu et al., 2016; Takahashi et al., 2005a; Takahashi et al., 2011), the auditory cortex exhibited organization into several auditory fields. The primary auditory cortex (A1) and the anterior auditory field (AAF) were characterized by short post-stimulus latency and showed a mirror image of distinct tonotopic gradients: A1 displayed a posterior-to-anterior gradient of low-to-high frequency in the dorsoposterior region, while AAF exhibited a high-to-low-frequency gradient in the ventro-anterior region (Fig. 2a, control). Other auditory fields showed longer latency than A1 and AAF and displayed tonotopic discontinuity in relation to the main tonotopic axes in A1 and AAF. The noise-exposed group exhibited a tendency to have fewer high-frequency regions (25–64 kHz) than those in the control group (Fig. 2a, exposed). This trend was further substantiated in the combined data analysis (Fig. 2b), where the proportion of recording sites with high CF significantly decreased (25–40 kHz, p=0.00109; 50–64 kHz, p=0.00264; two-tailed Mann–Whitney U test). In contrast, the proportion of recording sites with low-frequency CF significantly increased in the noise-exposed group (3.2–5.0 kHz: p=0.0162; 6.4–10 kHz: p=0.00295). Conversely, no significant difference was observed in the total number of recording sites with CF between the control and the noise-exposed groups (49.9±10.6 vs. 40.8±10.1; p= 0.0991, two-tailed Mann–Whitney U test).

**Figure 2.**
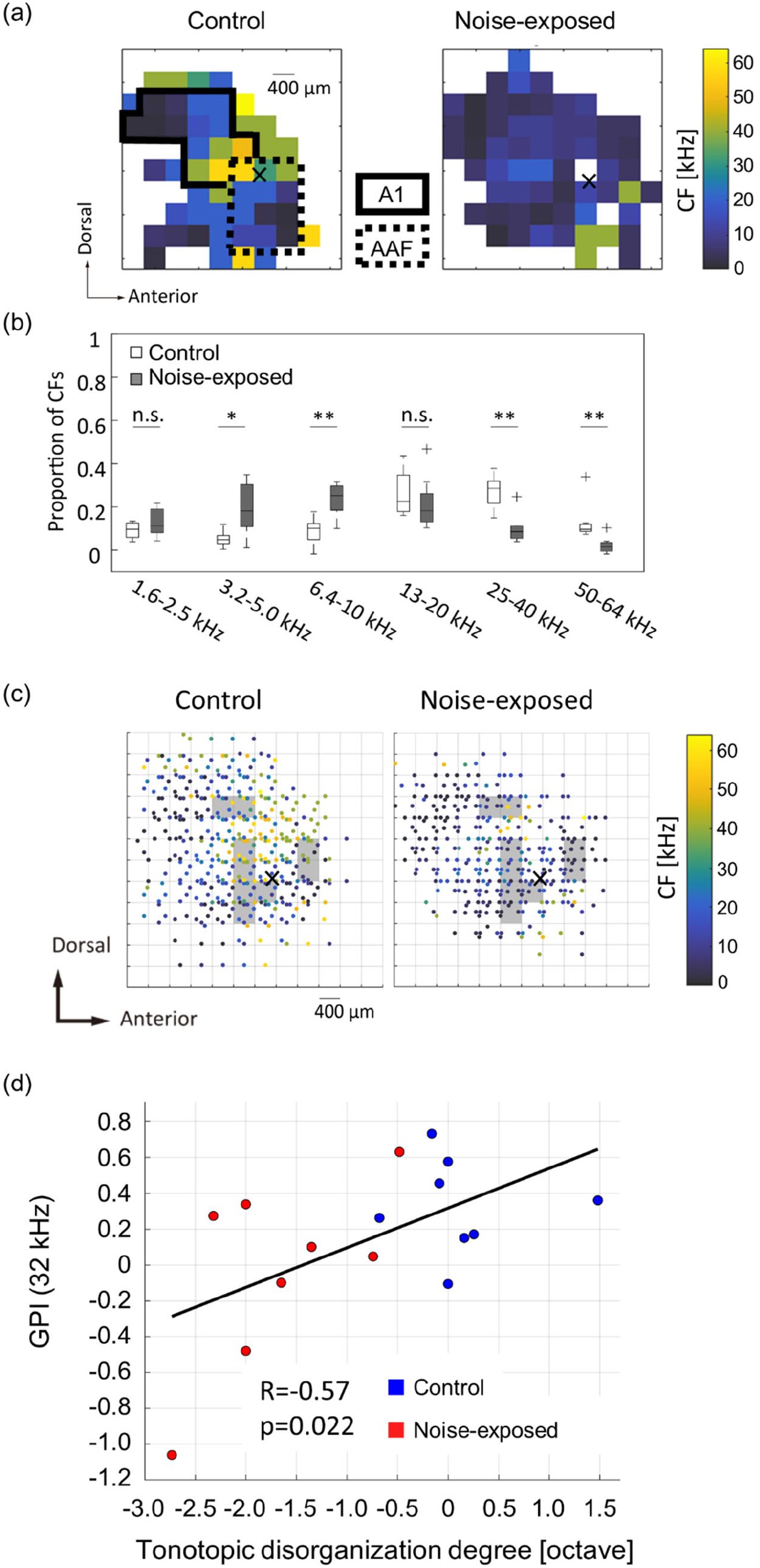
Acoustic trauma-induced disorganization represented in a tonotopic map. (a) Representative maps of characteristic frequency (CF) in the auditory cortex. Cross marks represent the activation focus of click-evoked LFP, which is used as a reference point to pool CF maps across animals. (b) Proportion of CFs in the control and exposed groups. Asterisks represent statistical difference between groups (*, p<0.05; **, p<0.01, Mann–Whitney U test). (c) Pooled CF maps. Statistical significances between the control and exposed groups were observed in shaded regions (p<0.05). (d) Correlation between the tonotopic disorganization degree and behavioral metrics of tinnitus (GPI).

**Figure 3.**
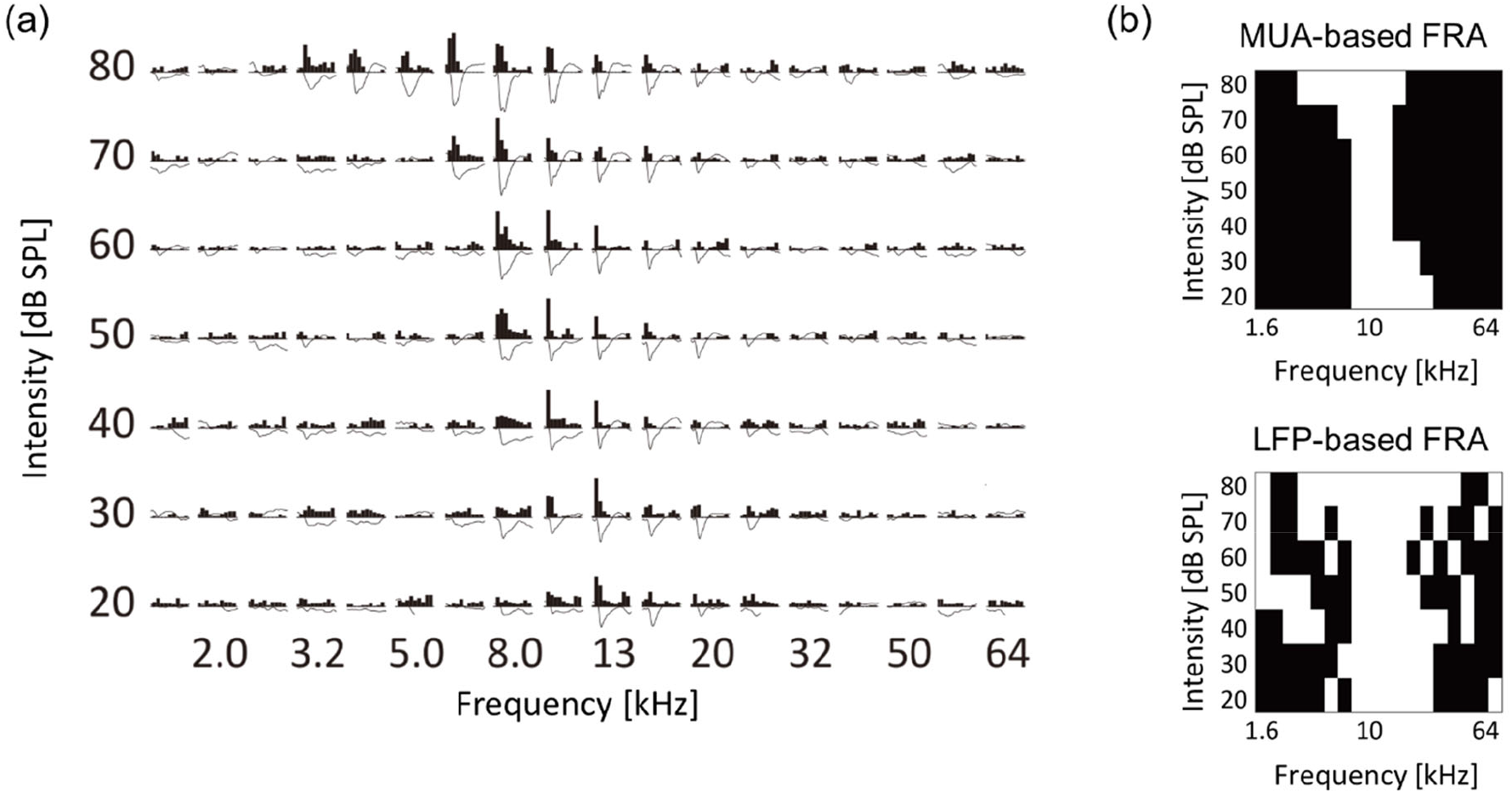
Characterization of tone-evoked neural activity. (a) Data from a representative recording site. Post-stimulus time histograms of MUA and stimulus-evoked LFP traces in response to test tones with different frequency and intensity are shown. MUA and LFP within the first 100 ms post-stimulus latency were characterized. (b) Binarized frequency response area (bFRA) of MUA and LFP. White colors represent activity (1) while black colors represent inactivity (0).

To assess acoustic trauma-induced disorganization of CF maps, CF maps from multiple animals were aligned and pooled using a reference point to create a group CF map (Fig. 2c). In accordance with our previous studies (Funamizu et al., 2013; Takahashi et al., 2005a; Takahashi et al., 2011), the reference point was determined at the activation focus of click-evoked LFPs by employing a quintic polynomial surface approximation for each animal (Figs. 2a and 2c, indicated by cross marks).

To identify the areas affected by noise exposure, CF maps were pooled on 400-μm grids and the intergroup differences were examined. Within each grid, corresponding to each square measuring 400 × 400 μm2, CFs values were gathered and compared between the groups. Significantly different CF values between the groups (two-tailed Mann–Whitney U test, p<0.05) were typically observed within CF regions ranging from 10 and 30 kHz (Fig. 2c, shaded areas), possibly corresponding to the “edge frequency” of the exposed tones. These shaded areas were regarded as the affected region.

To determine whether tonotopic disorganization within the auditory cortex is indicative of tinnitus symptoms (Adjamian et al., 2009; Engineer et al., 2011; Muhlnickel et al., 1998) or not (Elgoyhen et al., 2015; Koops et al., 2020; Langers et al., 2012), the correlation between the tonotopic disorganization degree and the behavioral index of tinnitus was explored. The tonotopic disorganization degree in each animal was defined as the deviation from the pooled CF map of the control group. In this evaluation, each animal’s CF within the affected region was compared to the median CF within a 200-μm radius of the corresponding test site on the pooled CF map of the control group. The median of these CF differences, expressed in octaves, was used to establish the degree of tonotopic disorganization for each animal, which was then plotted against the post-exposure GPIs at 32 kHz, serving as behavioral index of tinnitus at 32 kHz (Fig. 2d). The analysis revealed a significant correlation between the degree of tonotopic disorganization and the behavioral GPI index (R= 0.5679; t-test, p= 0.0217). In contrast, no significant correlation was observed between the tonotopic disorganization degree and the behavioral metrics of hyperacusis (PPI or ΔPPI) at any prepulse frequency (Supplementary Fig. S2). These results indicate that a pronounced disorganization of the tonotopic map is associated with severe tinnitus symptoms, but not with hyperacusis.

### 3.3 Cortical recruitment and neural gain

The examination focused on how the fourth layer of the auditory cortex was recruited for the neural representation of tones at the levels of MUA and LFP. These characteristics were analyzed for each test tone with varying frequencies and SPL (Fig. 3a). For each recording site, FRA values were calculated for MUA and LFP and then converted into binary values to classify whether a given recording site was active (1) or inactive (0) in response to each test stimulus (Fig. 3b). The threshold for binarizing FRA of MUA was determined as the inflection point of a smoothed z-score histogram of MUA(Guo et al., 2012; Noda and Takahashi, 2015). The threshold for the bFRA of LFP was set at 1.2 times the baseline fluctuations within 400–500 ms after the stimulus onset (Liu et al., 2015). Subsequently, by averaging bFRA across recording sites, the population bFRAs, or the CRF (Kilgard and Merzenich, 1998; Takahashi et al., 2011), were obtained for both MUA and LFP. These CRFs served as a measure of the activation level of the entire auditory cortex.

Figure 4 presents a comparison of group averages of CRFs at the MUA and LFP levels, denoted as CRF_MUA_ and CRF_LFP_, between the control and noise-exposed groups. The control auditory cortex demonstrated the highest recruitment in response to tones around 16 kHz (Fig. 4a, CRF_MUA_), whereas the auditory cortex in the noise-exposed group exhibited the largest recruitment to tones around 5 kHz and reduced recruitment to high-frequency tones compared to the control cortex (Fig. 4b, CRF_MUA_). Group comparison revealed significant effects of acoustic trauma: increased recruitment to low-frequency, high-SPL tones in CRF_MUA_ (white asterisks in Fig. 4b; p < 0.05, separate two-tailed Mann–Whitney U tests for each frequency and SPL level without multiple comparison adjustments) and decreased recruitment to high-frequency tones in CRF_MUA_ and CRF_LFP_ (black asterisks). Thus, the increase in CRF, a potential indicator of hyperacusis, was observed at the MUA level but not at the LFP level, suggesting that intracellular amplification from LFP to MUA at the thalamocortical recipient layer plays a critical role in hyperacusis. The acoustic trauma-induced CRF increases at moderate SPL in our results align with clinical observations that patients with hyperacusis perceive moderate-intensity sounds as intolerably loud, aversive, or painful (Anari et al., 1999; Auerbach et al., 2019; Auerbach et al., 2014; Baguley, 2003; Eggermont, J. J., 2017; Pienkowski, 2017, 2019).

**Figure 4.**
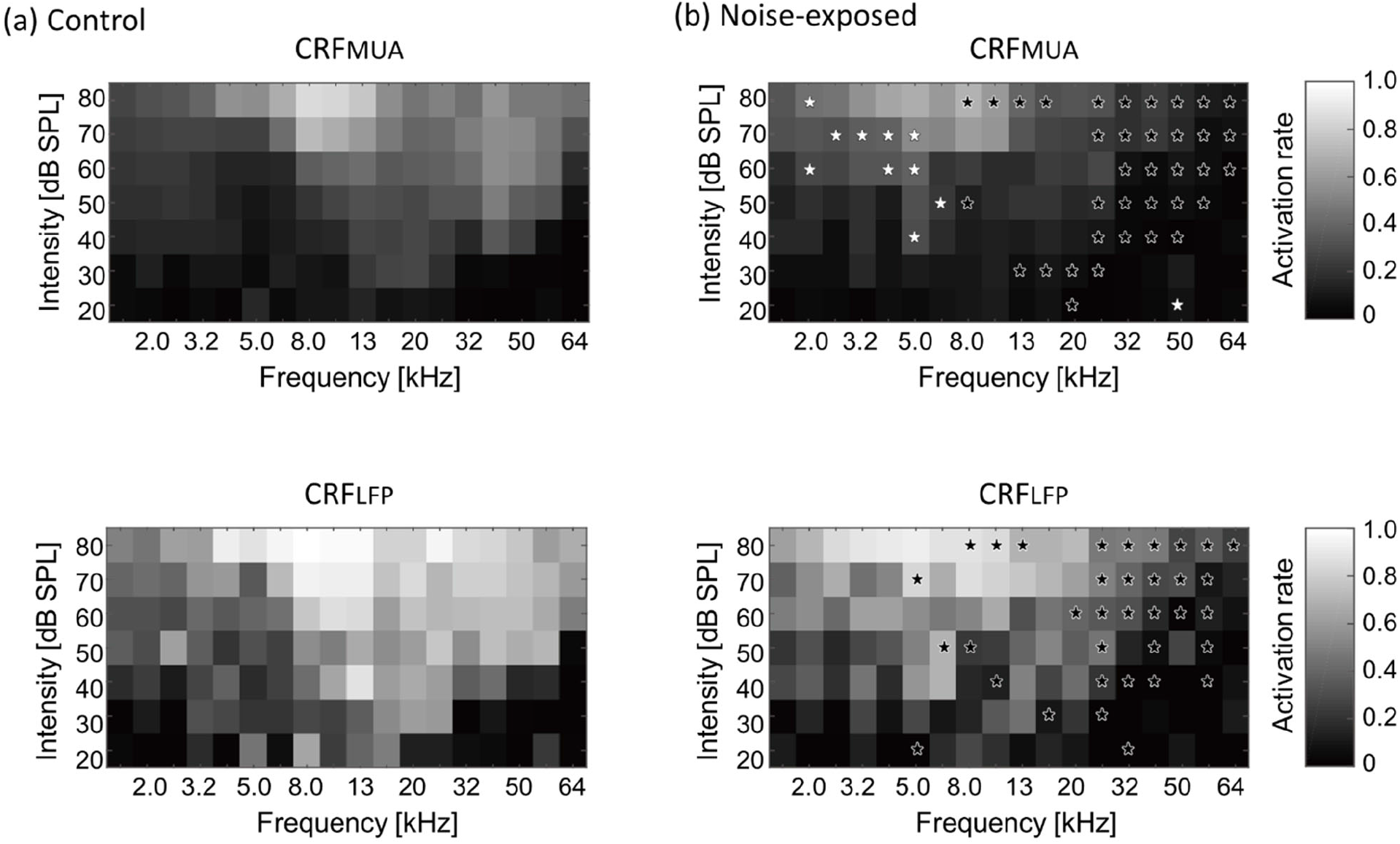
Cortical recruitment functions (CRFs) at the level of MUA and LFP (i.e., CRF_MUA_ and CRF_LFP_) in the control (a) and the exposed (b) groups. Black asterisks indicate decreased recruitment after the noise exposure, while white asterisks indicate increased recruitment (p<0.05, separate two-tailed Mann–Whitney U tests for each frequency and SPL level without multiple comparison adjustments).

To address the hypothesis that the above reorganizations of the map and CRF were associated with weakened lateral inhibition, which makes growth function of evoked activities against test intensity monotonic, the monotonicity index (MI) for each individual recording site was examined (Fig. 5a). The MI was defined as the ratio of the firing rate at the loudest SPL used (i.e., 80 dB SPL; red rectangles in Fig. 5a) to the maximum firing rate at the optimal SPL (i.e., blue rectangles in Figure 5a) (Zhou and Wang, 2010). MI takes a value ranging from 0 to 1, with higher values indicating greater monotonicity. Figure 5b characterizes MIs in either low (1.6–5.0 kHz), middle (6.4–20 kHz), or high (25–64 kHz) CF sites. MIs at the middle frequency region tended to show decreased proportion of moderately nonmonotonic neurons after noise exposure (Fig. 5c; 0.5<MI<1.0; Mann– Whitney U test, p=0.0531), exhibiting significant differences in distribution between pre- and post-exposure groups (Kolmogorov–Smirnov test: p=8.3e-04), suggesting that the lateral inhibition in the middle frequency weakened after the traumatic noise exposure. This weakened lateral inhibition may underlie the increased neural recruitment to low-frequency tones, which may be associated with hyperacusis symptoms.

**Figure 5.**
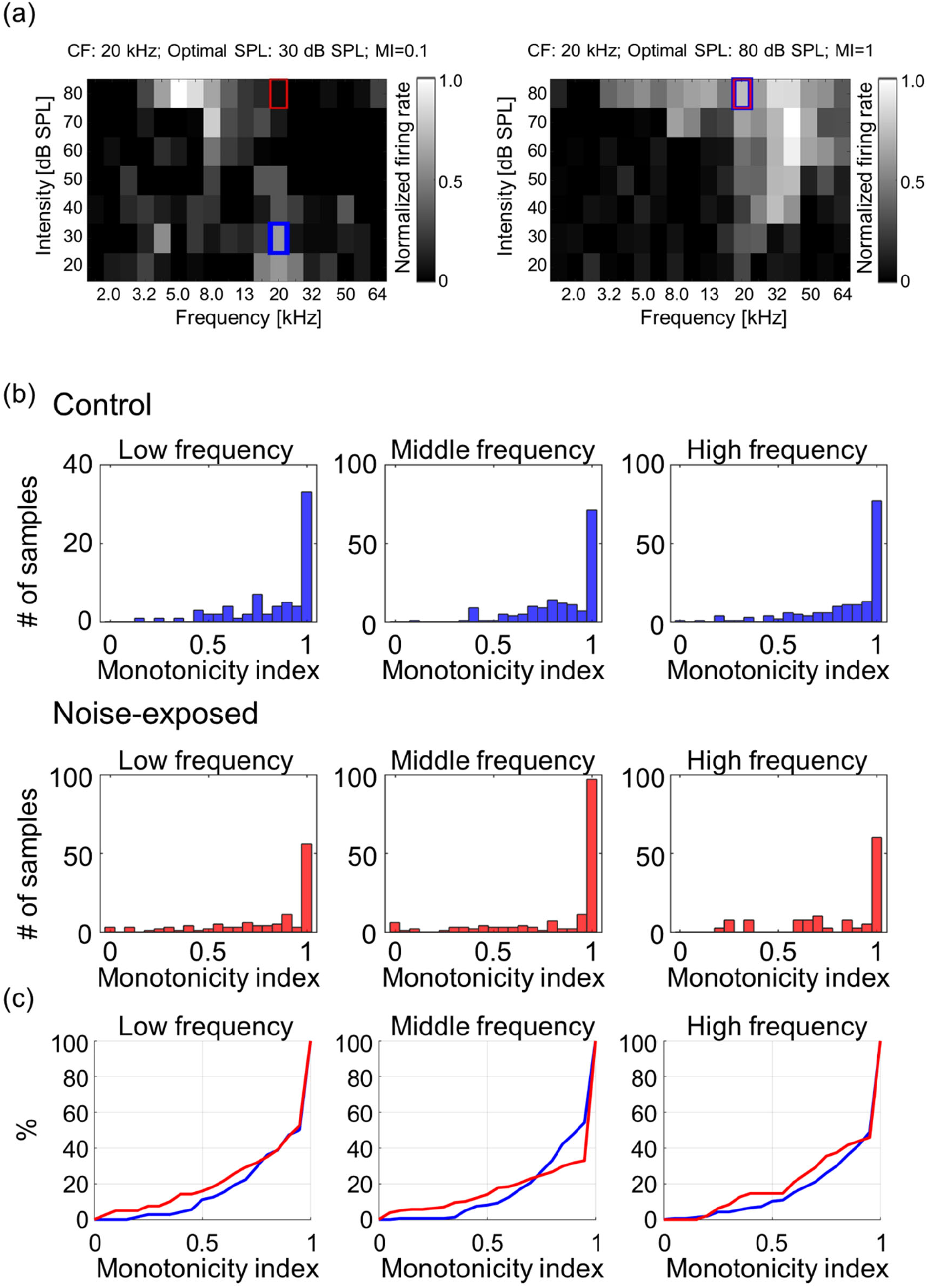
Monotonicity index (MI) of MUA. (a) Two examples of MI calculated as the ratio of the firing rate at the loudest SPL used (i.e., 80 dB SPL; red rectangles) to the maximum firing rate at the optimal SPL (i.e., blue rectangles). (b) Histograms of MI at low-, mid-, and high-frequency sites in the control and exposed groups. (c) Cumulative distribution of MI.

The increased recruitment at the level of MUA was hypothesized to contribute to hearing sensitivity in the noise-exposed animals and that the neurophysiological measure of CRF_MUA_ could predict the behavioral measure of hyperacusis, including ΔPPI at 4, 8, 16, and 32 kHz tones. To calculate CRF_MUA_ of each noise-exposed animal, the deviation from the group average of CRF_MUA_ in control animals was quantified and defined as ΔCRF_MUA_. At 4, 8, 16, and 32 kHz, the median of ΔCRF_MUA_ among 60–80 dB SPLs, or ΔCRF_MUA 60-80 dB SPL_ was quantified, since hyperacusis was commonly observed at moderate to high-SPL tones (Chen et al., 2014; Chen et al., 2015; Radziwon et al., 2019). Consequently, ΔCRF_MUA 60-80 dB SPL_ exhibited a significant positive correlation with ΔPPI (Fig. 6; R=- 0.39; t-test, p=0.038; y=1.56x+0.26), supporting our hypothesis. ΔCRF_MUA 60-80 dB SPL_ did not exhibited a significant correlation with the behavioral symptoms of tinnitus (ΔGPI at 32 kHz) (Supplementary Fig. S3; R=-0.3454, p=0.4480), suggesting that ΔCRF_MUA 60-80 dB SPL_ is neural correlates of hyperacusis, but not those of tinnitus.

**Figure 6.**
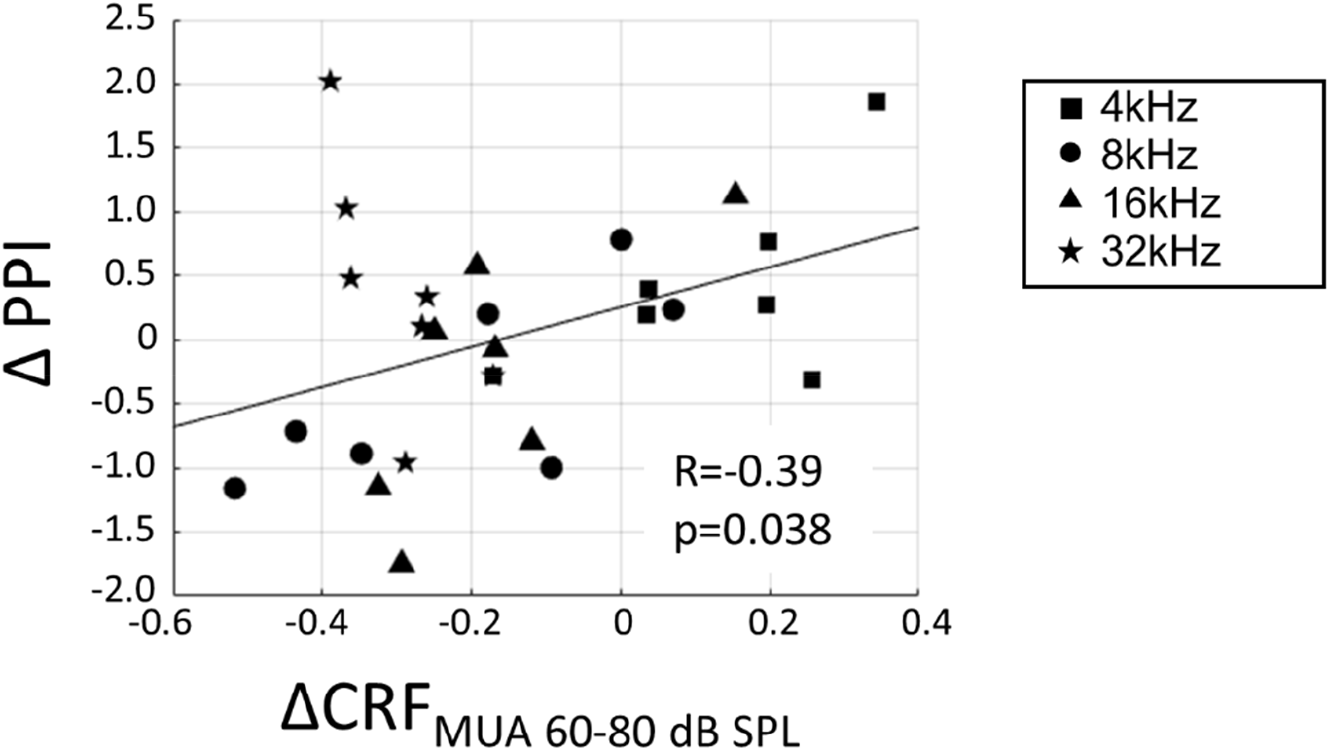
Correlation between hyperacusis symptoms (ΔPPI) and acoustic trauma-induced increase of CRF_MUA_ at moderate SPL (ΔCRF_MUA 60-80 dB SPL_). ΔPPI plotted against ΔCRF_MUA 60-80 dB SPL_ for tones at 4, 8, 16, and 32 kHz in the exposed group. A linear regression line is presented (*y*=1.56*x*+0.26).

As a measure of neural gain from LFP to MUA, the magnitude of MUA relative to LFP in the auditory cortex was characterized. Both LFP and MUA were normalized to each tone at each electrode concerning the maximum LFP and MUA (Fig. 7a), resulting in normalized LFP and MUA values ranging between 0 and 1. Then, for each animal, the ratio of MUA to LFP (MUA/LFP) for a tone frequency (f [kHz]) and SPL [dB SPL] was quantified as follows in Equation (2):

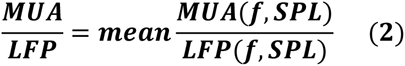

where ‘mean’ represented the average across electrodes that were active in the bFRA of LFP (Fig. 7b). Figure 7c illustrates MUA/LFP as a function of SPL by averaging MUA/LFP(f, SPL) across frequencies for each animal. While MUA/LFP decreased with SPL both the control and noise-exposed groups, indicating that the gain from LFP to MUA was high for low-SPL tones, this tendency was weaker in the noise-exposed group, especially below 60 dB SPL (Fig. 7c; p=0.048 for 30 dB SPL and p = 0.0499 for 40 dB SPL; Mann–Whitney U test without multiple-testing correction), suggesting that the dynamic range of the gain had narrowed in the noise-exposed group.

**Figure 7.**
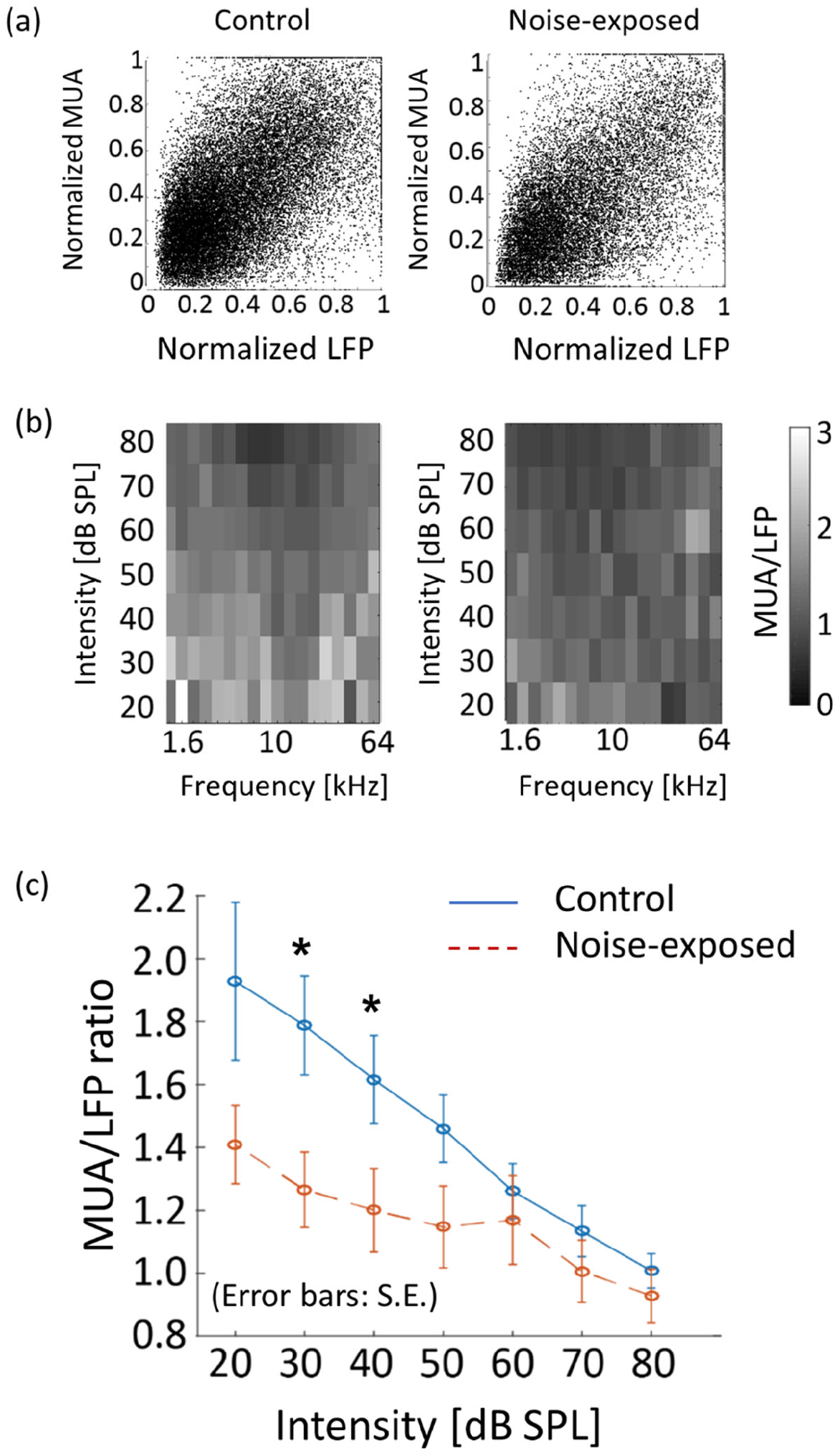
Neural gain from LFP to MUA (MUA/LFP) in the control and exposed groups. (a) Normalized MUA was plotted against normalized LFP. Each dot indicates a neural response to a tone. (b) MUA/LFP for each frequency-intensity condition. When the bFRA of LFP were inactive, MUA/LFP ratio was considered zero. (c) MUA/LFP as a function of intensity. Plots and error bars represent the average and errors. Asterisks represent statistical difference between groups (p<0.05, Mann–Whitney U test).

## 4 Discussion

We initially confirmed, through behavioral experiments, that rats exhibited different characteristics of PPI and GPI indices after being exposed to a traumatic 10-kHz tone. This observation suggests that the putative hyperacusis frequency differs from the tinnitus frequency. Although not reaching sufficient statistical levels due to wide variance across subjects, the U-shaped profile of ΔPPI indicated that the hearing impairment frequency was approximately between 8–16 kHz, while the possible hyperacusis frequency lay either below or above this hearing impairment frequency. Additionally, a reduction in GPI indicated that the subject had tinnitus in the high-frequency range, specifically approximately 32 kHz. Secondly, auditory cortex mapping revealed a significant correlation between GPI, a behavioral measure of tinnitus symptoms, and the extent of tonotopic map disorganization. Thirdly, ΔPPI, a behavioral index of hyperacusis symptoms, showed a correlation with the recruitment function at the MUA level in response to moderate- and high-SPL tones. However, this correlation was not observed at the LFP level. This suggests that hyperacusis was most prominent with high-SPL, low-frequency tones. The enhancement of MUA recruitment function was likely a result of increased gains in thalamocortical transmission, where LFP were considered as inputs and MUA as outputs. This gain modification was associated with the loss of monotonicity, a phenomenon in which inhibitory synapses played a crucial role.

### 4.1 Comparison with human studies

Our behavioral and electrophysiological findings suggest that exposure to a traumatic 10-kHz tone results in hearing loss with the 8–16 kHz range, tinnitus at a high frequency of approximately 32 kHz, and hyperacusis in response to moderate- and high-SPL tones at frequencies below 8 kHz. These findings align with human studies, where tinnitus pitch was most frequently observed at or above the frequency of the noise exposure (Atherley et al., 2005; Loeb and Smith, 2005; Sereda et al., 2011). However, it is worth noting that tinnitus frequencies have varied in previous animal models, sometimes falling below the noise exposure frequency (Engineer et al., 2011; Turner et al., 2006) or above it (Holt et al., 2010; Li et al., 2013; Longenecker and Galazyuk, 2011; Turner et al., 2012; Wang et al., 2009).

Nonetheless, our work may not be directly comparable to clinical studies. Firstly, human subjects with hyperacusis have shown reduced cortical activation in response to tones at the tinnitus frequency compared to those without hyperacusis (Koops and van Dijk, 2021). In our animal models, we did not observe a consistent trend between the putative tinnitus and hyperacusis frequency ranges. Secondly, the allocation of attention, wherein increased attention to hyperacusis frequency reduces attention to the tinnitus frequency, has been proposed as an underlying mechanism in the interaction between tinnitus and hyperacusis (Krumbholz et al., 2007; Paltoglou et al., 2011). However, our present study suggests an attention-free mechanism for tinnitus and hyperacusis. Our tonotopic map and cortical recruitment function, observed under anesthesia, demonstrated that acoustic trauma selectively reduced activation in high-frequency regions, including the tinnitus frequency, and conversely, induced hyperactivation in low-frequency regions at moderate and high intensities. Nevertheless, it is important to consider that acoustic startle responses are regulated by the reticular formation, i.e., the arousal system, which receives inputs from the auditory system and the amygdala (Carlson and Willott, 1998; Kandler and Herbert, 1991; Koch et al., 1992; Paus, 2000). Furthermore, the stress experienced by rats while being held in a confined chamber during GPI and PPI measurements may also affect their startle behavior (Eggermont, Jos J, 2017; Guercio et al., 2019). These emotional and conscious experiences should be taken into account. Therefore, further investigations are still required to validate our methods as translational studies of tinnitus and hyperacusis.

Clinically, noise-induced tinnitus can be acute or chronic. Acute tinnitus may last from a few minutes to several weeks after noise exposure(Han et al., 2009; Snow, 2004), while tinnitus that persists for several months is considered chronic(Mazurek et al., 2022) and tinnitus that persists for years is considered permanent and irreversible (Snow, 2004). Animal studies also showed that early signs of tinnitus and hyperacusis were sometimes reversible, while chronic signs developed over weeks (Hayes et al., 2014; Turner et al., 2012). Specifically, temporal elevation of auditory thresholds up to 30 dB was typically observed in the acute phase(Kujawa and Liberman, 2009; Middleton et al., 2011; Turner et al., 2012), which might be observed as the decreased PPI for BBN (Fig. 1a) and have confound effects in PPI and GPI interpretation. Therefore, our behavioral and electrophysiological experiments conducted shortly after noise exposure (2-8 days) are limited to the acute effects of noise exposure. Further studies are still required to investigate the neural correlates of chronic tinnitus and hyperacusis.

### 4.2 Neural correlates

Conceptual models of tinnitus and hyperacusis have been developed based on pioneering functional imaging studies in humans (Auer, 2008; Gu et al., 2010; Husain and Schmidt, 2014; Leaver et al., 2012; Llinas et al., 1999; Maudoux et al., 2012; Moazami-Goudarzi et al., 2010; Weisz et al., 2005). These studies have identified increased central gain, aberrant functional connectivity, and abnormal oscillations as the neural correlates (Henry et al., 2014; Sereda et al., 2011; Weisz et al., 2007). In addition to the group-level correlations between neural hyperactivity and hyperacusis-like behavior (Chen et al., 2014; Hickox and Liberman, 2014; Sun et al., 2009), we sought to demonstrate that examining inter-subject variability provides more robust evidence. This approach helps to illustrate how the loudness growth measured behaviorally is correlated with gain enhancement in the auditory cortex (Auerbach et al., 2019).

Our data suggests that tinnitus is linked to the disorganization of the tonotopic map. Tonotopic map disorganizations in subjects with tinnitus has been previously reported in humans (Adjamian et al., 2009; Muhlnickel et al., 1998) and in animals (Engineer et al., 2011), although these findings have remained controversial (Elgoyhen et al., 2015; Koops et al., 2020; Langers et al., 2012). We believe that these discrepancies arise because human studies on tonotopic mapping often used moderate-intensity tones to evoke distinct cortical activation, while in animal studies, CFs in tonotopic maps were defined based on the lowest intensity tones that activated the test neuron. Our data align with previous research indicating that the hyperactivity induced by salicylate and trauma in the auditory cortex is associated with a FRA shift toward the mid-frequency region (Chen et al., 2014; Chen et al., 2012; Chen et al., 2013; Norena et al., 2010; Stolzberg et al., 2011). The central noise hypothesis of tinnitus is consistent with the disorganization of the tonotopic map characterized with low-intensity tones, which evoke responses with relatively low S/N ratios. In this context, map plasticity characterized by low-intensity tones may be the potential neural correlates of tinnitus but not hyperacusis.

The perceived loudness depends on both the intensity and the bandwidth of the test stimulus, which is related to the number of activated frequency channels (Gu et al., 2010; Hawley et al., 2005; Zwicker et al., 1957). Therefore, the recruitment function, which quantifies the number of activated neurons in response to a given test tone, serves as a suitable predictor of how loudly the test tone is perceived. Neural recruitment typically corresponds to the activation of a neural population. Activation of the primary auditory cortex increases with intensity in both individuals with normal and impaired hearing (Behler and Uppenkamp, 2016; Hall et al., 2001; Langers et al., 2007). Likewise, hyperacusis, in both human and animals, is commonly associated with hyperactivity in higher-order subcortical nuclei and the auditory cortex (Auerbach et al., 2019; Chen et al., 2015; Gu et al., 2010; Harms and Melcher, 2002; Knipper et al., 2013; Ruttiger et al., 2013; Sigalovsky and Melcher, 2006; Zeng, 2013), rather than with broadening of cortical tuning (Koops and van Dijk, 2021). This hyperactivity has been observed over a wide range of test frequencies beyond regions of hearing loss (Diehl and Schaette, 2015; Noreña and Chery-Croze, 2007; Sheldrake et al., 2015). Additionally, earplugging and acoustic enhancements also induce adaptive gain control mechanisms across frequency channels (Formby et al., 2003; Munro et al., 2014; Noreña and Chery-Croze, 2007).

Our results indicate that hearing loss led to reduced recruitment of neural populations in response to high-frequency tones, but increased recruitment in response to high-SPL, low-frequency tones, specifically at the MUA level, but not at the LFP level (Fig. 4). Since the first negative deflection of LFP reflects synaptic inputs to the thalamocortical layer (Einevoll et al., 2013; Haider et al., 2016; Mazzoni et al., 2015), while MUA reflects outputs resulting from nonlinear intracellular processing in the cortex, our findings suggest that the neural correlate of hyperacusis is associated with the output rather than the input of the thalamocortical input layer. Consequently, hyperacusis was likely to manifest in response to high-SPL, low-frequency tones, which is consistent with previous research indicating that sound-evoked hyperactivity is not limited to frequencies affected by hearing loss in patients with hyperacusis (Koops and van Dijk, 2021) and in animal models (McGill et al., 2022). Figure 6 also supported our hypothesis in that the sign of hyperacusis (ΔPPI) was correlated with increased recruitment of MUA to high SPL tones (ΔCRF_MUA 60-80 dB SPL_); however, this was true for 4-16 kHz tones, but not for 32 kHz tone. Map plasticity was also frequency dependent, increasing in low CF regions and decreasing in high CF regions (Fig. 2). These data suggest that additional CF-dependent mechanisms, which cannot be captured by ΔCRF_MUA 60-80 dB SPL_, underlie the compensatory gain control and behavioral symptom of hyperacusis.

We showed that comparison of CRF between LFP and MUA served as a possible measure to characterize how efficiently the thalamo-cortical synaptic inputs (LFP) were converted into cortical discharges (MUA), i.e., the neural gain from the LFP to MUA. Thalamocortical transmissions in the noise-exposed group were characterized by narrower dynamic ranges of input/output ratios compared to those in the control group (Fig. 7). These results align with previous studies demonstrating that cochlear damage reduces sound-evoked neural responses at the auditory nerve (Heinz et al., 2005; Heinz and Young, 2004; Wake et al., 1993), leading to an enhanced gain, characterized by a steep increase in neural response with sound intensity, at the level of the auditory cortex (Asokan et al., 2018; Chambers et al., 2016; Jiang et al., 2017; McGill et al., 2022; Norena et al., 2003; Parameshwarappa et al., 2022; Popelar et al., 1987; Qiu et al., 2000; Resnik and Polley, 2017; Resnik and Polley, 2021; Seki and Eggermont, 2003; Syka et al., 1994).

Our analyses were made possible because both auditory-evoked LFP and MUA were distinct at layer 4 in the auditory cortex. However, we cannot definitively conclude that our findings are specific to the thalamocortical layer. Similar analyses could be employed to investigate whether and how the gain from synaptic inputs to neuronal discharges varies among auditory subcortical nuclei and different layers of the auditory cortex following hearing loss. In previous studies, tinnitus and hyperacusis have been linked to increased auditory-evoked LFP and fMRI responses in the IC, MGB and A1. These findings were considered evidence of central gain enhancement following hearing loss, possibly through homeostatic plasticity (Auerbach et al., 2014; Chen et al., 2015; Gu et al., 2010; Qiu et al., 2000; Salvi et al., 1990). Central noise, thought to underlie tinnitus but not hyperacusis, may also increase with central gain in some models (Norena, 2011) but not in others (Zeng, 2013). Due to these discrepancies, hyperacusis may not always be associated with tinnitus (Baguley, 2003). The hyperactivity in hyperacusis gradually develops through the auditory brainstem pathway in a sequential manner and is most consistently observed at the level of the auditory cortex (Chen et al., 2015; Qiu et al., 2000; Schaette and McAlpine, 2011). These cumulative effects may result from nonlinear processing from LFP to MUA at each nucleus, as demonstrated at the level of the thalamocortical recipient layer in our current study.

### 4.3 Neural mechanisms

The gain control mechanisms underlying tinnitus and hyperacusis are likely triggered by partial impairment of the peripheral auditory pathway. For instance, rats exhibited clearer evidence of tinnitus after exposure to noise at 110 dB SPL compared to exposure at 116 dB SPL or higher (Turner and Larsen, 2016). In mice, spontaneous activities in the IC increased more following a 2-hour noise exposure at 100 dB SPL than at 105 dB SPL. In our study, we utilized a unilateral hearing loss model, known for its effectiveness in inducing tinnitus and hyperacusis (Isaacson and Vora, 2003; Jahn and Polley, 2023). This model may be accompanied by neural plasticity at various auditory processing centers, including the cochlear nucleus (Rubio, 2006), lateral superior olive (Kotak and Sanes, 1995), IC (Vale and Sanes, 2000), and the auditory cortex (Kotak et al., 2008; Sarro et al., 2008). Severe bilateral hearing loss could deprive the higher-order auditory system of effective gain control.

The changes in gain are associated with increased gene expression of glutamate receptors and decreased expression of GABA receptors in the auditory cortex (Balaram et al., 2019; Sarro et al., 2008). This suggests hypersensitization and disinhibition, respectively. Particularly, parvalbumin-expressing interneurons play a crucial role in triggering the hyperactivity of cortical pyramidal neurons (Masri et al., 2021; Resnik and Polley, 2017; Resnik and Polley, 2021).

We have demonstrated that hearing loss significantly reduces the monotonicity of tone-evoked activities, suggesting that a loss of inhibition underlies the modulation of neural recruitment induced by acoustic trauma. Auditory-evoked potentials in the primary auditory cortex exhibit a linear response to the rate of pressure change (in Pa/s) when stimulated with CF tone but a nonlinear response to non-CF tones. This suggests that inhibition plays a role in the nonlinearity characteristics of loudness perception (Takahashi et al., 2004, 2005b). The loss of inhibition, or disinhibition, is the most likely mechanism behind central gain enhancement (Auerbach et al., 2014; Chen et al., 2014; Norena, 2011). Disinhibition resulting from acoustic trauma aligns with findings that acoustic trauma reduces tuning to CF tones (Scholl and Wehr, 2008) and broadens the FRA in auditory cortex neurons, extending far beyond the CF (Salvi et al., 2000; Wang et al., 1996). This suggests that individual neurons receive a wide range of inhibition. Similarly, high-dose treatment with sodium salicylate induces tinnitus and hyperacusis and enhances gain in the central auditory system (Auerbach et al., 2019; Auerbach et al., 2014; Hayes et al., 2014; Jiang et al., 2017), most notably in the auditory cortex (Norena et al., 2010; Yang et al., 2007; Zhang et al., 2011), along with hyperactivity in non-auditory systems (Chen et al., 2014; Chen et al., 2016; Chen et al., 2017; Chen et al., 2015).

Both noise trauma and salicylate treatment are likely to result in synaptic disinhibition (Lu et al., 2011; Milbrandt et al., 2000; Sun et al., 2009; Wang et al., 2011; Yang et al., 2011). Salicylate suppresses GABA-mediated inhibition and enhances excitability (Gong et al., 2008; Xu et al., 2005), and systemic salicylate application induces hyperactivity in the auditory cortex, leading to symptoms of tinnitus and hyperacusis (Chen et al., 2012; Sun et al., 2009), which can be ameliorated by enhancing GABA-mediated inhibition (Brozoski et al., 2007; Lu et al., 2011; Sun et al., 2009). The compensatory plasticity resulting from the loss of inhibition is already observed at the level of the cochlear nucleus (Fang et al., 2016; Ngodup et al., 2015; Salvi et al., 2000). Our results align with the observation that hyperacusis is often experienced over a broad frequency range, including lower frequencies with normal hearing thresholds (Anari et al., 1999; Noreña and Chery-Croze, 2007; Sheldrake et al., 2015). These properties of hyperacusis are attributed to the loss of lateral inhibition, which has been observed from the region of central hearing loss to lower frequency regions (Auerbach et al., 2014). Without lateral inhibition, frequencies near the lesion edge could become “over-represented” in the central auditory system, leading to the recruitment of excess neurons in the auditory cortex (Eggermont, J. J., 2017). Our findings suggest that the negative effects of the loss of lateral inhibition are more pronounced at moderate and high intensities than at lower intensities. It is important to note that the neural activities in our study were characterized under anesthesia, which could have significant effects on neural gain and sound-evoked activities (Noda and Takahashi, 2015; Ros et al., 2017; Szalda and Burkard, 2005; Thornton and Sharpe, 1998; Yang et al., 2007). Since isoflurane increases inhibitory tones, the effects of disinhibition may be more pronounced in awake conditions than those characterized in our study.

Our findings were based on the collective activities of neurons. Imaging with cellular resolution provided additional insights, revealing that changes in gain following acoustic trauma varied among pyramidal neurons at cortical layer 2/3. Specifically, the gain remained stable in pyramidal neurons with low spontaneous activity and nonmonotonic intensity tuning, likely due to strong inhibition (Tan et al., 2007; Wu et al., 2006). Conversely, non-homeostatic gain control was typically observed in neurons with high spontaneous activity and monotonic tuning (McGill et al., 2022). At the synaptic level, the balance between excitatory and inhibitory inputs underlies the frequency tuning of auditory cortical neurons (Wehr and Zador, 2003; Zhang et al., 2003). The loss of inhibition may lead to CF upshifts in low-CF neurons and CF downshifts in high-CF neurons (Scholl et al., 2008; Wang et al., 2002). These observations are consistent with the map plasticity induced by acoustic trauma in the auditory cortex, which includes an expanded representation in the 3.2–6.4 kHz regions and reduced representation in the 25–40 kHz regions.

Tinnitus and hyperacusis are likely to co-occur, possibly because the auditory cortex plays a central role in the tinnitus-hyperacusis network, which includes connections with the amygdala, the reticular formation, the hippocampus, striatum, and the cerebellum (Carlson and Willott, 1998; Chen et al., 2014; Chen et al., 2015; De Ridder et al., 2006; Gu et al., 2010; Hayes et al., 2014; Paus, 2000; Salvi et al., 2021; Ulanovsky and Moss, 2008; Zeng, 2013). This interconnected network may lead to a dissociation between cortical and subcortical neural activities (Boyen et al., 2014). Tinnitus is associated with altered functional networks extending beyond the auditory system (Chen et al., 2015; Husain and Schmidt, 2014; Leaver et al., 2012; Llinas et al., 1999). Specifically, increased functional connectivity between the auditory cortex and the amygdala, along with hyperactivity in the amygdala, are commonly observed in both patients with tinnitus and animal models (Aazh et al., 2018; Chen et al., 2012; Chen et al., 2016; Chen et al., 2013; Hazell and Jastreboff, 1990; Jüris et al., 2013; Kim et al., 2012; van Veen et al., 1998; Wallhäusser-Franke et al., 2003). This plasticity is likely driven by homeostatic plasticity, through which the auditory system overcompensates for the reduced output at the peripheral cochlear level (Auerbach et al., 2014; Norena, 2011).

In conclusion, our study in the auditory cortex has revealed a correlation between the disorganization of the tonotopic map and tinnitus-like symptoms, as well as an association between increased unit activity and hyperacusis. These findings indicate that neural correlates of tinnitus can be identified in population firing responses to low-intensity tones, while those of hyperacusis are linked to high-intensity tones. These results support the central noise hypothesis in tinnitus and the maladaptive gain control hypothesis in hyperacusis. To the best of our knowledge, this study is the first to disentangle the neural correlates of tinnitus and hyperacusis within the auditory cortex. We believe that our research offers a novel perspective on the neural foundations of tinnitus and hyperacusis resulting from noise-induced hearing loss.

## 7 Author Contributions

NW: Data curation, Formal Analysis, Funding acquisition, Investigation, Methodology, Visualization, Writing – original draft

TIS: Funding acquisition, Methodology, Validation, Writing – review & editing

HT: Conceptualization, Funding acquisition, Project administration, Supervision, Writing – review & editing

## 9 Data availability

Data supporting the conclusion of this article will be made available on a reasonable request.

## Supplementary Material

### 1 Supplementary Figures

**Supplementary Figure S1.**
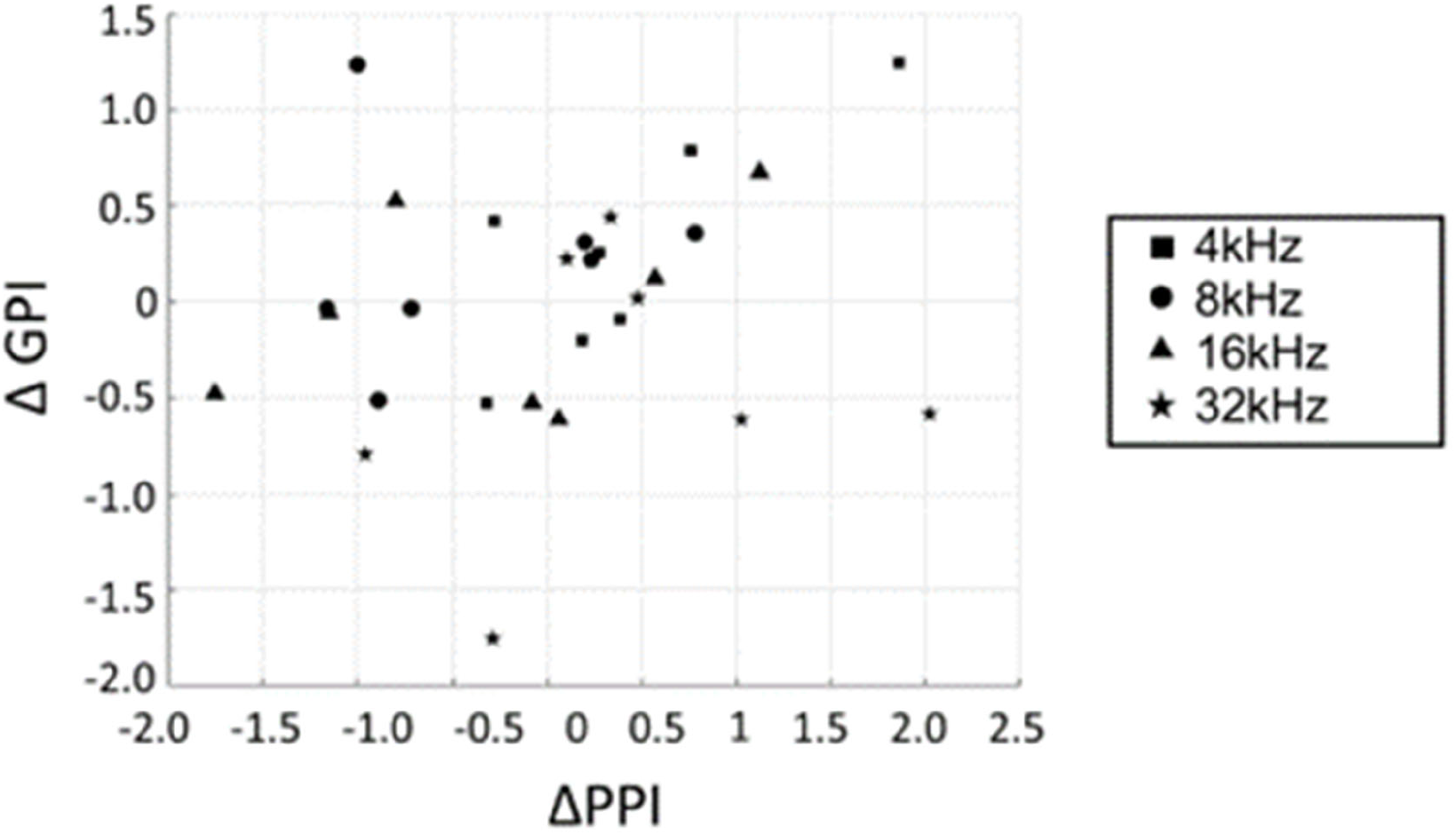
Behavioral metrics of hyperacusis (ΔPPI) vs. tinnitus (ΔGPI).

**Supplementary Figure S2.**
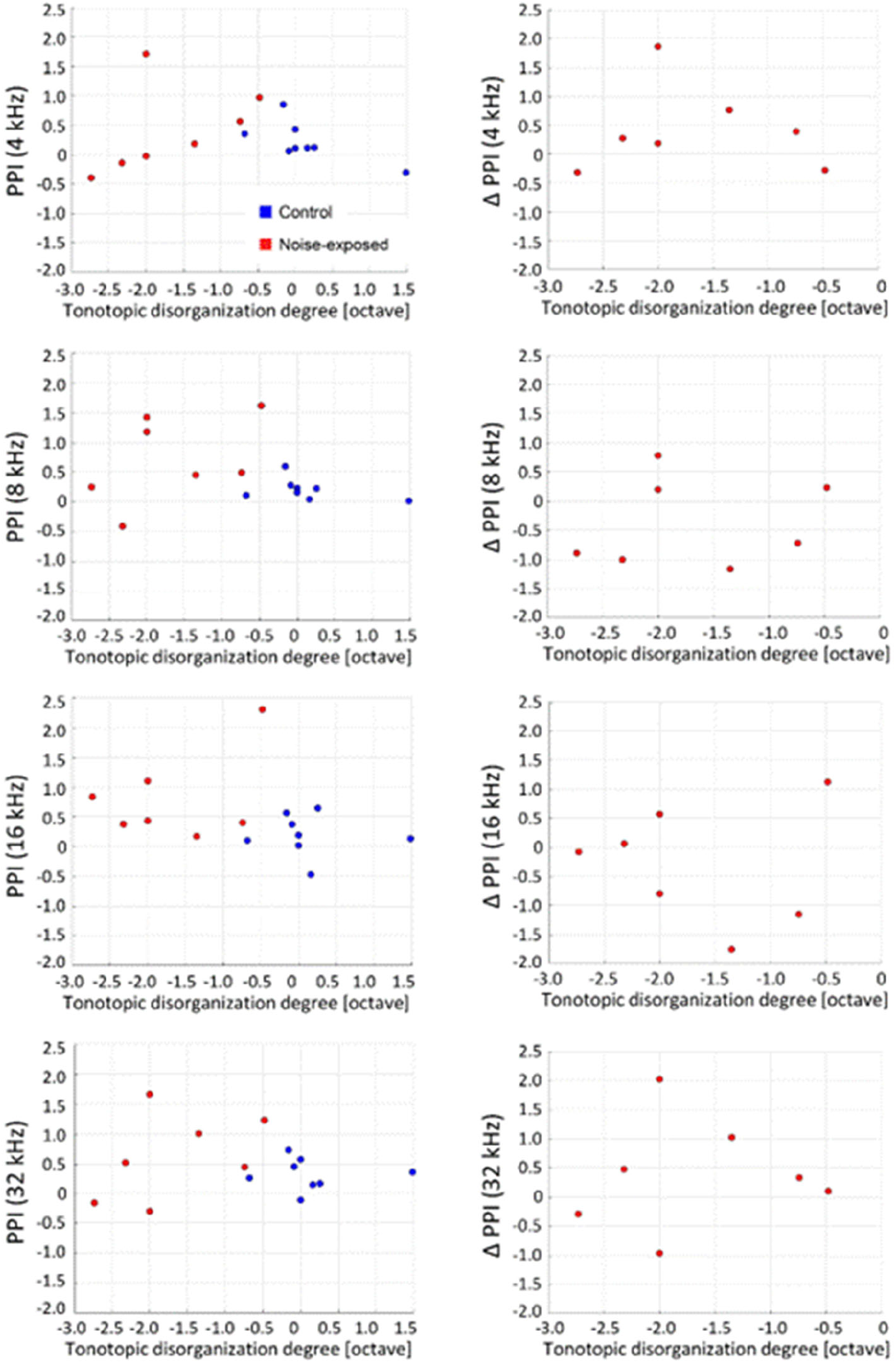
Behavioral metrics of hyperacusis (PPI and ΔPPI) plotted against the tonotopic disorganization degree in the exposed group. Correlation coefficients in PPI with respect to the tonotopic disorganization degree (left column): R=-0.0684 (t-test, p=0.809) at 4 kHz; R=-0.229 (p=0.411) at 8 kHz; R=-0.265 (p=0.341) at 16 kHz; R=-0.0655, p=0.817 at 32 kHz. ΔPPI (right): R=- 0.0727 (t-test, p=0.877) at 4 kHz; R=0.143 (p=0.760) at 8 kHz; R=-0.0314 (p=0.947); and R=0.0803 (p=0.864) at 32 kHz.

**Supplementary Figure S3.**
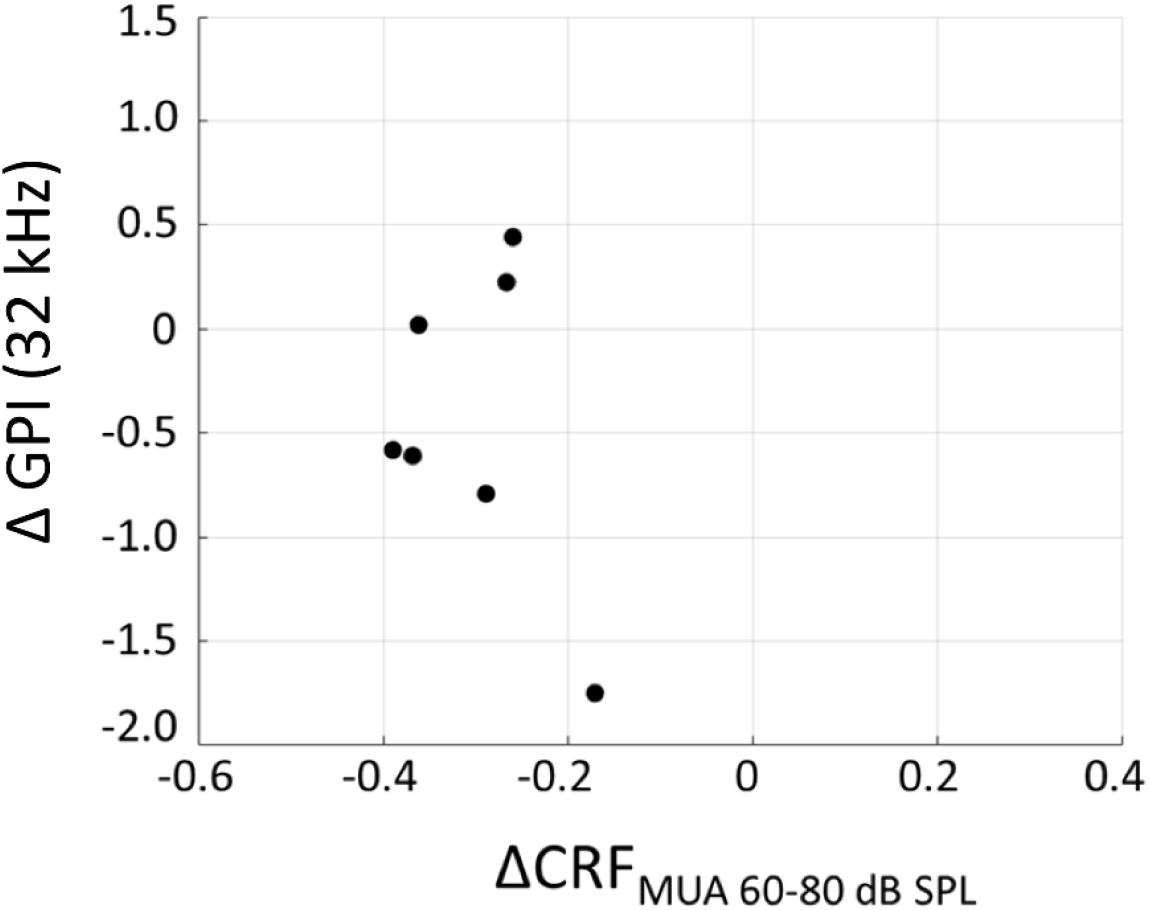
Behavioral metrics of tinnitus (ΔGPI at 32 kHz) plotted against ΔCRFMUA 60-80 dB SPL. Correlation coefficients: R=-0.3454 (t-test, p=0.4480).

